# The Time-Course of Food Representation in the Human Brain

**DOI:** 10.1101/2023.06.06.543985

**Authors:** Denise Moerel, James Psihoyos, Thomas A. Carlson

## Abstract

Humans make decisions about food every day. The visual system provides important information that forms a basis for these food decisions. Although previous research has focused on visual object and category representations in the brain, it is still unclear how visually presented food is encoded by the brain. Here, we investigate the time-course of food representations in the brain. We used time-resolved multivariate analyses of electroencephalography (EEG) data, obtained from human participants (both sexes), to determine which food features are represented in the brain, and whether focused attention is needed for this. We recorded EEG while participants engaged in one of two tasks. In one task the stimuli were task relevant, whereas in the other task the stimuli were not task relevant. Our findings indicate that the brain can differentiate between food and non-food items from approximately 112 milliseconds after stimulus onset. The neural signal at later latencies contained information about food naturalness, how much the food was transformed, as well as the perceived caloric content. This information was present regardless of the task. Information about whether food is immediately ready to eat, however, was only present when the food was task relevant and presented at a slow presentation rate. Furthermore, the recorded brain activity correlated with the behavioural responses in an odd-item-out task. The fast representation of these food features, along with the finding that this information is used to guide food categorisation decision-making, suggests that these features are important dimensions along which the representation of foods is organised.

## 1. INTRODUCTION

Humans make food choices every day. Our visual system allows us to gather information about distant foods, making visual information an important basis for food choices. There is evidence that food processing through vision and taste rely on overlapping processes, with activation in the same regions (Simmons et al., 2005), and shared information about the taste (Avery et al., 2021), evoked by food that is seen and tasted (cf. Nakamura et al. (2020)). Recent evidence suggests that food could be an important organisational feature in the ventral visual stream, as neural responses in the ventral visual cortex are selective for visual food stimuli (Jain et al., 2023; Khosla et al., 2022). Previous work has shown that information about animacy, category, and exemplars emerges in the brain before 100 ms (Carlson et al., 2013; Cichy et al., 2014; Contini et al., 2017; Grootswagers et al., 2019). However, relatively few studies have focused on the time-course of visually presented food representations.

Several studies have investigated the neural food/non-food distinction in the temporal domain (Stingl et al., 2010; Tsourides et al., 2016). For example, one multivariate magnetoencephalography (MEG) study showed that evidence for information about food vs. non-food emerges from 85 ms after stimulus onset (Tsourides et al., 2016). Fewer studies have investigated whether, and when, other features of food are represented in the brain. Two dimensions that are of relevance from an evolutionary perspective are 1) the distinction between natural and prepared food (i.e., food with signs of human intervention), and 2) the energetic value of the food (i.e., calorie content).

The distinction between natural and prepared food may be an important one for modern humans. Even Chimpanzees have been shown to spontaneously prefer cooked food (Warneken & Rosati, 2015; Wobber et al., 2008). Several studies have investigated the neural distinction between natural and prepared foods (Coricelli et al., 2019; Pergola et al., 2017; Vignando et al., 2019), suggesting that the neural response is modulated by whether food is natural or prepared. However, it is still unclear whether the brain carries information about this distinction, and what the time-course of this information is.

Another important feature of food is the energetic value or caloric content. Several (electroencephalography) EEG and functional Magnetic Resonance Imaging (fMRI) studies have found that the brain response to food stimuli is modulated by the caloric content of the stimuli (Killgore et al., 2003; Mengotti et al., 2019; Meule et al., 2013; Toepel et al., 2009) or perceived healthiness of the food (Rosenblatt et al., 2018). However, it remains unclear whether, and at what time, the brain carries information about the energetic content of food.

In addition to features such as naturalness and calorie content, the way the brain processes food information could depend on perceiver characteristics. For example, one study used multivariate pattern analysis on EEG data to show there was information about the subjective ‘tastiness’ of the food from approximately 530 ms after stimulus onset (Schubert et al., 2021). It is still unclear whether the brain represents the level of arousal in addition to the valence (palatability) of the food. In addition, it is unclear whether general food preferences across the population can be used to track valence information in the brain, or whether more fine-grained personal valence information is needed.

Here, we used multivariate analysis methods on EEG data to investigate the time-course of edibility information, naturalness, calories, perceived immediate edibility, valence and arousal information in the brain. In addition, as previous MEG work showed effects with short latencies and in the absence of a task (Tsourides et al., 2016), we asked whether these representations emerge even when the stimuli are not the focus of attention. Finally, to determine whether the food feature information is represented in a way that is accessible to the brain, and used to guide behaviour, we correlated the EEG representations to behavioural food similarity judgments from a different group of participants.

## 2. MATERIALS AND METHODS

### 2.1. Participants

Twenty healthy adults participated in the EEG study. The participants had a mean age of 20.9 years (SD = 3.74, range 18-33 years). Seventeen participants identified as female, two as male and one as non-binary. All participants were right-handed, reported normal or corrected-to-normal vision, and had no history of psychiatric or neurological disorders. The participants were recruited from the University of Sydney and received course credit for their participation. The study was approved by the Human Ethics Committee of the University of Sydney (ethics ID: 2019/340) and all participants provided informed consent before participating.

### 2.2. Experimental design

The stimulus set consisted of 154 food images and 160 non-food object images (Figure 1A). The food images were taken from the Foodcast Research Image Database (Foroni et al., 2013). The subset was chosen to be familiar to Australian participants, and no rotten food items were included. We selected 71 natural food items and 83 transformed (or prepared) food items. The non-food stimuli were from a free image hosting website (PNG Imaging) and were a subset of those used in previous EEG object decoding studies (Grootswagers et al., 2019). The non-food stimulus set consisted of 80 animate and 80 inanimate objects. In this study, we define animacy as whether the object can move on its own volition, and it is therefore different than aliveness. We used the animacy distinction, as this has been extensively used in previous work (Carlson et al., 2013; Cichy et al., 2014; Contini et al., 2017; Grootswagers et al., 2019). The 160 non-food object stimuli were divided into 8 categories (humans, mammals, birds, insects, plants, clothing, furniture, and tools). Each category had 5 different objects. For example, the ‘mammal’ category consisted of cow, giraffe, etc. There were four exemplars of each object. Note that the fruit category from this set was not included because they are foods. The aquatic category was also excluded because of some exemplars might be seen as unprepared foods (e.g., fish or squid).

**Figure 1.**
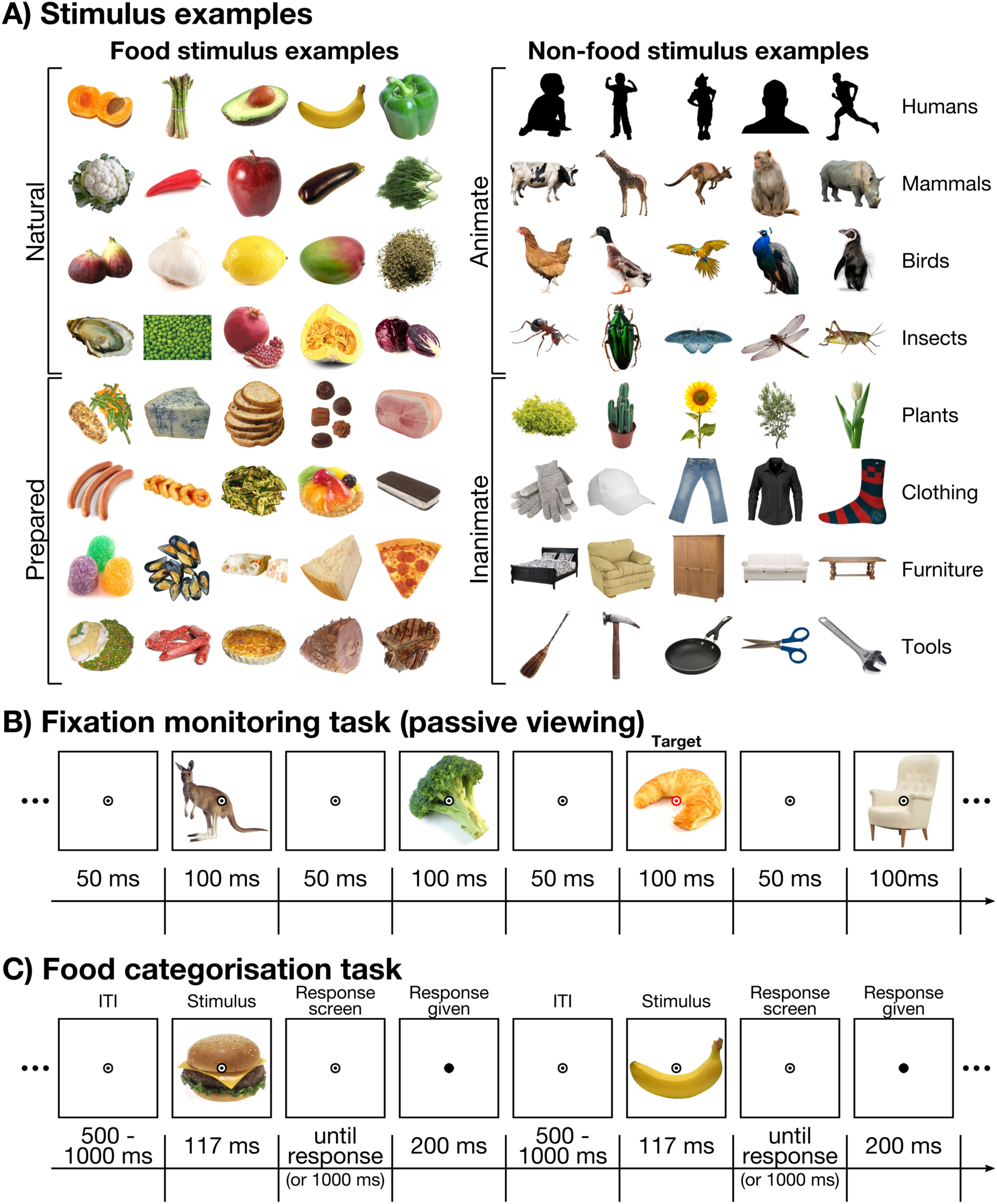
Stimuli and task. **A)** Shows examples of the stimuli. The left panel shows examples of the 154 food stimuli that were used. Stimuli were selected from the Foodcast Research Image Database (Foroni et al., 2013). The selected stimulus set consisted of 71 natural food images (top) and 83 transformed food images (bottom). The right panel shows examples from the 160 non-food object stimuli. Images were obtained from a free image hosting website (PNG Imaging) and were used in previous work (Grootswagers et al., 2019). The stimulus included 80 animate (top) and 80 inanimate objects (object) from 8 different categories (humans, mammals, birds, insects, plants, clothing, furniture, and tools). Note the stimuli depicting humans (top row) are replaced with silhouettes in this figure to comply with bioRxiv guidelines. **B)** Shows an overview of the fixation monitoring task, in which the stimuli were not task relevant. Stimuli were presented for 100 ms with an inter-stimulus-interval of 50 ms. Participants pressed a button when the fixation bullseye changed from black to red. The third stimulus on this example (croissant) shows an example of a target. **C)** Shows an overview of the food categorisation task. Participants saw a food stimulus for 117 ms and were asked to indicate whether the depicted food item was natural or prepared. There was a response time-out of 1000 ms. When a response was given, the fixation bullseye turned into a black dot for 200 ms. There was a randomly selected inter-trial-interval ranging between 500 ms and 1000 ms.

We used the Psychopy library (Peirce et al., 2019) in Python to run the experiment. The images were presented on a white background and were 256 by 256 pixels. Participants were seated approximately 60 cm from the screen, and the stimuli were approximately 6.25 by 6.25 degrees of visual angle. A central fixation bullseye was presented throughout the experiment, and participants were instructed to remain fixation on the central bullseye. The EEG experiment consisted of two different tasks, a fixation monitoring task and a food categorisation task (Figure 1B and 1C). There were 6 blocks in total, and each block consisted of the fixation monitoring task followed by the food categorisation task. The experiment took approximately 50 minutes to complete. Participants used a button box to respond in both tasks.

#### 2.2.1. Fixation monitoring task (passive viewing)

Figure 1B gives an overview of the fixation monitoring task. During this task, participants viewed streams of images. The images were presented at 6.67 Hz; each stimulus was on the screen for 100 ms, followed by a blank screen for 50 ms. The stimuli consisted of 314 unique images, depicting food (154 images) and non-food objects (160 images). There were 6 blocks in the experiment. Each block consisted of 6 image sequences, and each sequence contained 157 image presentations. This means there were 942 image presentations per block, 3 repeats of the 314 unique images. The order of the stimulus presentations was randomised across two consecutive sequences. Each sequence was approximately 24 seconds, and participants were given the option to have a break after every sequence to minimise fatigue and eye blinks.

The participants monitored the fixation bullseye and pressed a button when the fixation bullseye turned to red instead of black for the duration of a stimulus presentation (100 ms). Participants were instructed to respond as fast and accurately as possible. The purpose of the task was to keep participants engaged and ensure their gaze remained fixated on the centre of the screen. Each sequence had between 2 and 4 targets (fixation colour changes). The targets were presented at random intervals with the following constraints. The first ten and last ten stimulus presentation of the sequence were never targets, and two targets were always at least 15 stimulus presentations apart.

#### 2.2.2. Food categorisation task

Figure 1C provides an overview of the food categorisation task. In this task, participants indicated for each image whether the depicted food was ‘natural’ or ‘prepared’. The naturalness distinction was chosen as a task because previous work suggests that naturalness could be an important feature in how we process food. Several studies have found evidence the neural response is modulated by whether food is natural or prepared (Coricelli et al., 2019; Pergola et al., 2017; Vignando et al., 2019). In addition, this distinction could be important for behaviour, as there is evidence that naturalness mediates the behavioural ratings of other food features such as arousal and perceived calorie content (Foroni et al., 2013). The stimuli were the same as those used in the fixation monitoring task, but only the food images were used. Each stimulus was presented once per block in a randomised order. Participants completed 6 blocks in total. The food stimulus was presented for 117 ms, and participants had 1000 ms to make a response. If participants responded, the fixation bullseye turned to a filled black dot for 200 ms. This was done to indicate to the participant that their response was recorded. If no response was made within 1000 ms, feedback was given that the response was too slow, and the experiment automatically moved on to the next trial. There was an inter-trial interval ranging between 500 ms and 1000 ms. Participants were instructed to respond as fast and accurately as possible. They responded by pressing one of two buttons on a button box. The buttons used to indicate whether the food was ‘natural’ or ‘prepared’ switched between each block. This was done to make sure that any differences in the neural signal between natural and prepared foods could not be driven by motor preparation or execution. At the start of each block, instructions were shown detailing the button-response mapping for that block. Participants were given the opportunity to take a break after every 22 trials to avoid fatigue.

### 2.3. EEG acquisition and pre-processing

The EEG data were recorded with a 128 channel BrainVision ActiChamp system, at a sampling rate of 1000 Hz, referenced online to FCz. The electrodes were placed according to the international 10-20 system (Oostenveld & Praamstra, 2001). We used a minimal EEG pre- processing pipeline, following earlier work (Grootswagers et al., 2021; Robinson et al., 2021). First, we re-referenced to the average of all electrodes. Then we applied a low-pass filter of 100 Hz and a high pass filter of 0.1 Hz, and we down-sampled the data to 250 Hz. Finally, we created epochs locked to each stimulus onset, from -100 ms before stimulus onset to 1200 ms after stimulus onset.

### 2.4. Decoding analysis

We used decoding analyses to determine *if* and *when* there was information in the EEG data about two dimensions: 1) whether an object is a food or non-food item, and 2) whether a food is natural or prepared. For these analyses, we used a linear discriminant analysis to distinguish the classes of interest on the pattern of activation across all electrodes, for each individual time-point, and for each individual participant, using the CoSMoMVPA toolbox for Matlab (Oosterhof et al., 2016). The classifiers were trained and tested on data from single trials, no trial averaging was used. For the EEG data obtained during the fixation monitoring task, we used an exemplar-by-block cross validation approach (Carlson et al., 2013; Grootswagers et al., 2019). We used a pair of images, one from each class, from one block as test data. We trained the classifiers on the remaining images from the remaining 5 blocks in an iterative way. Images were paired up at random. We repeated this analysis 10 times with different random image pairings and averaged across the obtained decoding accuracies. Each of the random image pairing iterations ran through the full cross-validation process, leaving each block out as a test block once. We used exemplar-by-two-blocks cross validation procedure for the data obtained during the food categorisation task. Participants used button presses to indicate whether a food was natural or prepared during this task, which means that motor preparation or execution could contaminate the signal for this task. We left out two blocks with opposite response button mapping as test data, iterating over all 9 possible combinations of two test blocks with opposite button mapping.

To determine the time-course of the representation of edibility, we trained the classifier to distinguish food from non-food objects for each time point. This analysis was done for the fixation monitoring task only because non-food objects were not included in the food categorisation task condition. We balanced the groups of food and non-food items, making sure there were an equal number of food and non-food stimuli. To determine the time-course of food naturalness representations, we trained the classifier to distinguish between natural and prepared food. We did this for both the fixation monitoring task and the food categorisation task. For the fixation monitoring task, only the food items were included. We balanced the groups of natural and prepared stimuli. As a comparison for the time-course of edibility decoding, we also decoded animacy for the fixation monitoring task. Only the non-food items were selected for this analysis. To match the power between the animacy and edibility analyses, we repeated the edibility decoding analysis matching the number of training trials with the animacy analysis. Decoding analyses were done separately for each individual participant, and Bayes factors were calculated at the group level.

Previous work has shown that eye movements could be a possible confounding factor in decoding analyses (Quax et al., 2019). The fixation bullseye was presented during the stimulus presentation, as well as between stimulus presentations, and participants were instructed to remain fixated on the bullseye. The rapid (100 ms) stimulus presentation in both tasks removes the incentive to make eye movements. In addition, the next stimulus is presented 150 ms after the previous one in the fixation monitoring task, which means the next stimulus will already be on the screen once the saccade is completed. Although the fast timing of both tasks mitigates the contribution of eye moments to the decoding accuracy, we cannot fully rule out their contribution to the decoding, as all electrodes were used in the analyses described above. We anticipated that posterior regions would drive the decoding of visual information in this experiment, whereas a potential contribution of eye movements to the signal would be expected in frontal electrodes that are located near the eyes (Lins et al., 1993). To determine whether posterior brain regions were indeed driving the decoding, we repeated the same decoding analyses described above using a subset of the following 56 posterior electrodes: CP1, CP2, CP3, CP4, CP5, CP6, CPP1h, CPP2h, CPP3h, CPP4h, CPP5h, CPP6h, CPz, O1, O10, O2, O9, OI1h, OI2h, Oz, P1, P10, P2, P3, P4, P5, P6, P7, P8, P9, PO10, PO3, PO4, PO7, PO8, PO9, POO1, POO10h, POO2, POO9h, POz, PPO10h, PPO1h, PPO2h, PPO5h, PPO6h, PPO9h, Pz, TP10, TP7, TP8, TP9, TPP10h, TPP7h, TPP8h, TPP9h.

This analysis follows the logic of previous work, using a 64-channel set-up, where the posterior 28 electrodes were selected (Robinson et al., 2021). Although the contribution of eye movement signals cannot be fully ruled out, this method uses data that is unlikely to be contaminated by eye movements.

### 2.5. Representational similarity analysis

To gain insight into what features of foods are represented in visual processing dynamics, we used Representational similarity analysis (RSA) (Kriegeskorte et al., 2008; Kriegeskorte & Kievit, 2013). RSA Representational Dissimilarity Matrices (RDMs) encode the dissimilarity between two food items. We obtained an RDM based on the EEG data (see Figure 2A) for each time-point, individual participant, and for the two different tasks (fixation monitoring and food categorisation). In the fixation monitoring task, we used only the food stimuli to construct the RDMs. Pair-wise decoding was used as a measure of neural dissimilarity, computed using the same decoding methods described above. We made seven models each describing a feature of the foods: naturalness, perceived transformation, perceived immediate edibility, real calories, perceived calories, valence, and arousal (Figure 2B). All models were computed based on data included in FRIDa (Foroni et al., 2013), and therefore based on data from a different group of participants. The naturalness and real calorie models were based on objective values, whereas the other models were based on the subjective behavioural ratings of 73 participants. The naturalness model coded for the dissimilarity in whether the food was natural (i.e., whether it can be found in nature) or prepared (see Figure 3A). The perceived transformation model coded for the dissimilarity in how much a food was perceived to have been transformed from its natural state (i.e., how processed the food is) (see Figure 3B). Therefore, the differences between the naturalness and perceived transformation models are 1) the naturalness model is objective whereas the perceived transformation model reflects subjective ratings from participants, and 2) the naturalness model is binary, whereas the perceived transformation model is continuous. For example, in the naturalness model, both pickles and lasagne are classified as prepared foods. However, in the perceived transformation model, lasagne is rated as more transformed than pickles. The perceived immediate edibility coded for the dissimilarity in the amount of time and effort that was perceived to be required before the food could be eaten (see Figure 3B). Both natural and prepared foods can be immediately edible, and perceived immediate edibility is therefore not related to naturalness or perceived transformation. For example, an apple or cupcake is immediately edible, whereas dried beans or uncooked pasta require time to prepare. The real calorie model was based on the calorie content of 100g of the food. The perceived calorie model was based on subjective behavioural ratings of the caloric content of 100g of the food. The valence model was based on how positive or negative the item represented in the image was rated. Finally, the arousal model represents the emotional salience of the items, as rated by the participants. Because the valence and arousal model were based on data from a different group of participants, they do not capture the personal preferences of the EEG participants. Rather, if there is a correlation between the EEG and these models, this will provide evidence for the coding of information about preference that generalise across the population. For each time-point and individual EEG participant, we correlated the EEG RDM with each of the seven food feature models.

**Figure 2.**
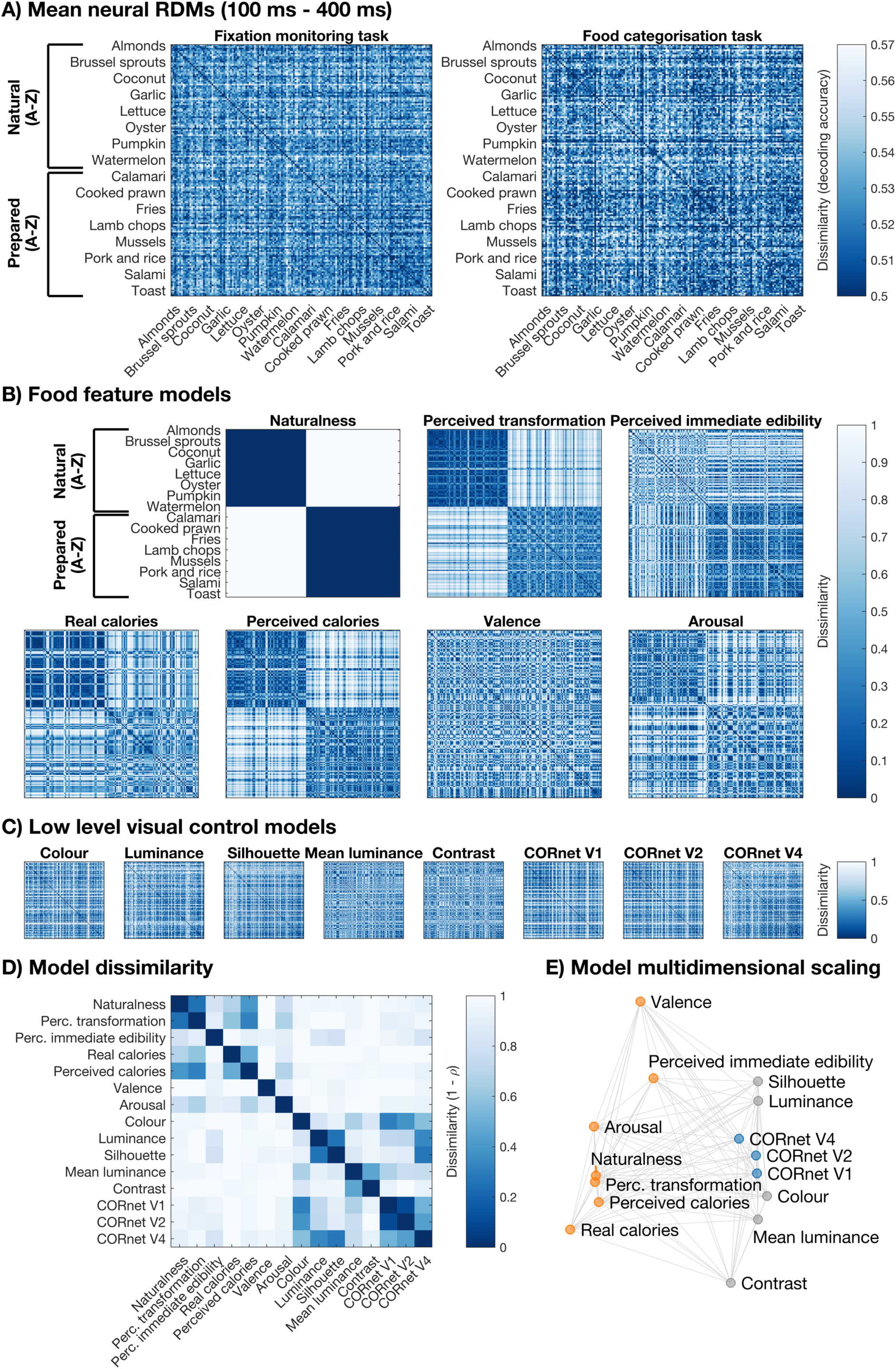
Representational similarity analysis overview. **A)** Shows the representational dissimilarity matrix (RDM) for the EEG data for the fixation monitoring task (left) and the food categorisation task (right), averaged over a 100 ms to 400 ms time-window for visualisation. The RDM is a 154 by 154 matrix and each point codes for the dissimilarity between two food stimuli. The food items are ordered as follows: all natural foods in alphabetical order, followed by all prepared foods in alphabetical order. **B)** Shows the seven food feature model RDMs. All points in the RDMs refer to the same combinations of food items as A. The top row, from left to right, shows the naturalness model, perceived transformation model, and perceived immediate edibility model. The bottom row, from left to right, shows the real calorie model, perceived calorie model, valence model and arousal model. **C)** Shows the five low-level image feature control model RDMs and three CORnet-S deep convolutional artificial neural network models. We used these models control for visual differences between the groups of food stimuli. **D)** Shows the dissimilarity between all models: the seven food feature models, five low-level image feature control models, and three CORnet control models. The dissimilarity is calculated as 1 – the correlation (Spearman’s *ρ*). **E)** Shows a 2-dimensional projection of all model dissimilarities. The distance between two models approximates their dissimilarity. The configuration was obtained through multi-dimensional scaling. The food feature models are shown in orange, the low-level image feature control models in grey, and the CORnet control models in blue.

**Figure 3.**
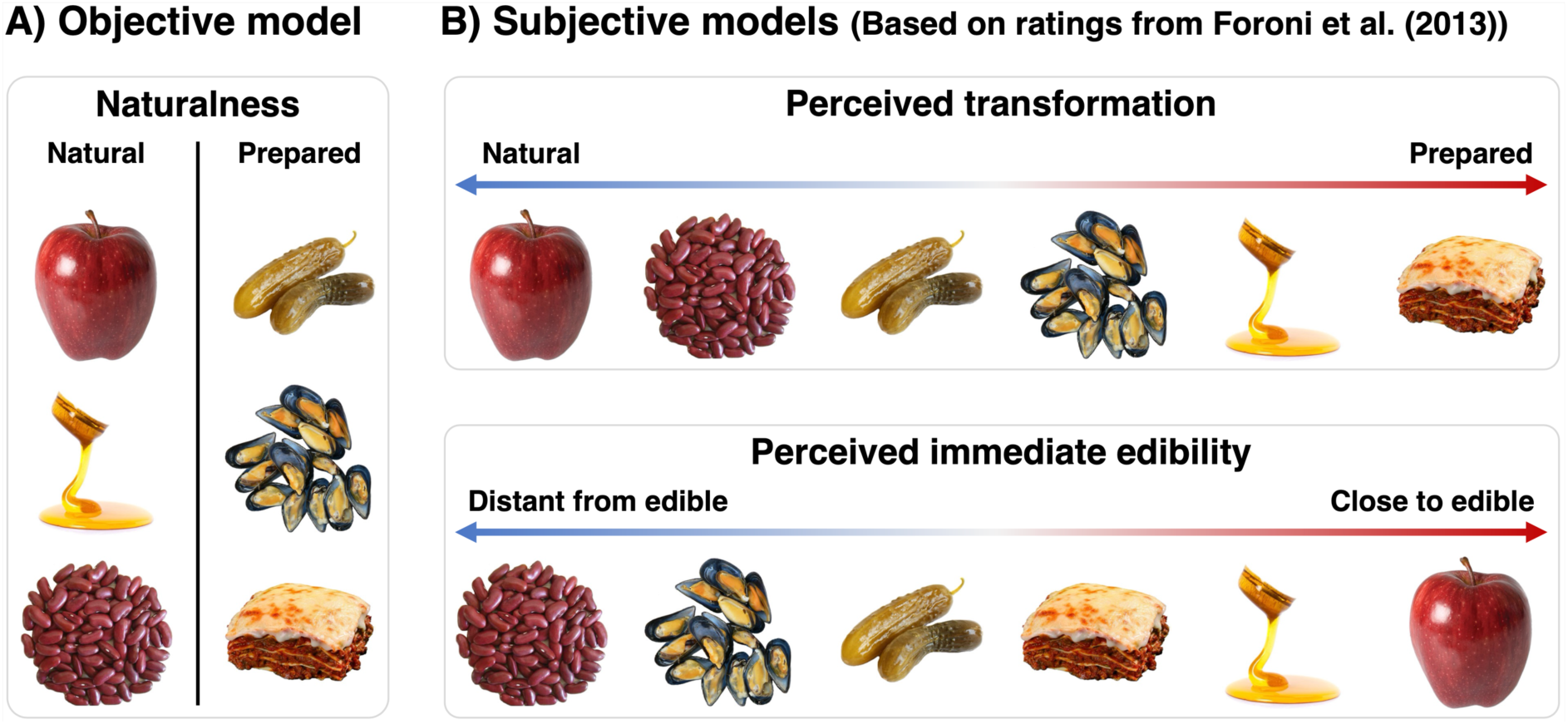
Food feature model examples. **A)** shows and example of stimuli arranged according to the naturalness model. This model is based on objective categorisation of whether a food is natural or prepared. This model is binary, where a food item is either classified as natural or prepared. **B)** shows examples of stimuli arranged according to two subjective models, that are based on ratings from a different group of participants (Foroni et al., 2013). These models are continuous. The top panel shows the perceived transformation model, with stimuli perceived to be more natural on the left and those perceived to have undergone more preparation on the right. The bottom panel shows the perceived immediate edibility model. Foods that are perceived to require the most intervention before they can be consumed are shown on the left, and foods that are immediately ready to eat are shown on the right.

To account for low-level visual differences between the stimulus groups, we partialled out control models from the correlations between the EEG and food models (Figure 2C). There were three models based on deep neural networks (CORnet-S) and five low-level image feature control models. We used the *‘partialcorr’* Matlab function from the Descriptive Statistics toolbox to calculate the partial rank correlations between the EEG and food feature model RDMs, controlling for the low-level visual and CORnet control models. Figure 2D shows the dissimilarity (1-correlation) between the seven food feature models, the five low-level image feature control models, and the three deep neural network models, and Figure 2E shows the multidimensional scaling of the model dissimilarities. The five low-level image feature control models included a silhouette model, a pixel-wise colour model, a pixel-wise luminance model, a mean luminance model and an RMS contrast model. The control models were based on the colour/luminance values of the images and were not measured on the experiment screen. The silhouette model, pixel-wise colour model, and pixel-wise luminance models were based on vectorised images. The silhouette model used the Jaccard distance between the binary alpha layer of each pair of images as a measure for dissimilarity. The pixel-wise colour model and pixel-wise luminance models were based on the CIELAB colour values and greyscale values respectively, using correlation as a distance measure. We used the rgb2lab function in Matlab to convert from RGB to CIELAB, assuming that the white point was D65 and the colour space of the screen was sRGB. The mean luminance model and RMS contrast model were based on a value calculated across the entire image (Harrison, 2022), using the mean greyscale values of the stimuli and the standard deviation of the greyscale values of the stimuli respectively, and the mean difference as a distance measure. In addition, we used CORnet-S (Kubilius et al., 2019), a deep convolutional artificial neural network, to control for low-level visual differences between the stimulus classes. To obtain the models, we used the openly available CORnet-S code (https://github.com/dicarlolab/CORnet) on the images from this experiment. We used the layers “V1”, “V2”, and “V4”, designed to be analogous to the ventral visual areas V1, V2, and V4 respectively. We then used a cosine distance metric to obtain the dissimilarity for each pair of stimuli in the RDM.

To determine if information about whether an object is a food could be driven by low-level visual differences between the stimulus groups, we made EEG RDMs for all stimulus items, food and non-food, presented in the fixation monitoring task. We correlated this to an edibility model that coded for the dissimilarity in whether an item was a food or non-food object. We ran partial correlations in two separate analyses, partialling out 1) the five low-level image feature control models and 2) the three CORnet-S control models described above.

It is important to note that the naturalness, perceived transformation, and perceived calorie models are correlated. It is possible that *naturalness* is driving the correlation between these models. The naturalness model provides an objective and binary description of naturalness (natural or prepared), whereas the perceived transformation model is based on continuous subjective ratings on the naturalness/transformation dimension. There is evidence that the perceived calorie model also has information about naturalness, as the perceived calorie ratings are biased by whether the item is natural or prepared. Participants were more likely to underestimate the calories in natural foods and overestimate the calories in prepared foods (Foroni et al., 2013). This means that it is possible that naturalness is the underlying dimension linking these models. We examined which of these correlated food feature models explain unique variance in the EEG data. We used partial correlations to examine the contribution of the naturalness, perceived transformation, and perceived calorie models. To examine the contribution of the naturalness model, we calculated the full correlation between the EEG RDM and the naturalness model, as well as three partial correlations, partialling out 1) the perceived transformation model, 2) the perceived calorie model, and 3) both. We followed the same logic for the perceived transformation model. Here, we calculated the full correlation with the EEG, and again three partial correlations, partialling out 1) the naturalness model, 2) the perceived calorie model, and 3) both. Finally, to assess the contribution of the perceived calorie model, we calculated the full EEG correlation and four partial correlations, partialling out 1) the naturalness model, 2) the perceived transformation model, 3) the real calorie model, and 4) all of these models.

### 2.6. Investigating the link between brain and behaviour

The analyses described above can provide insight into which features of food stimuli are represented in the brain. There is often an implicit underlying assumption that above chance decoding accuracy or EEG-model correlation means this information is represented in a way that is accessible by the brain. However, evidence of information alone is not enough to make this claim (de-Wit et al., 2016). One way to determine whether these food features are represented in an accessible way, is to determine whether they are used by the brain to guide behaviour. To establish a link between the brain’s representation and behaviour, we ran an online odd-one-out triplet experiment (Hebart et al., 2020) with new participants. In this experiment, participants were presented with three food images on the screen and asked to select the odd-one-out, using any criteria. The advantage of the triplet odd-item-out task is that the third food acts as a context for other two foods, allowing us to determine which features are used to make a decision. This allows us to establish a behavioural model of human food discrimination that encompasses multiple features. We constructed a behavioural similarity model, where similarity was measured as the probability of two foods being chosen together (i.e., if two foods were presented together, what was the probability that the third food was selected as the odd-one-out). We then correlated the behavioural similarity model to the EEG RDM, separately for each individual participant, time-point, and for the two different EEG tasks (fixation monitoring and food categorisation). By recruiting a different group of participants for the behavioural and EEG experiments, we can assess a general link between the coding of food features in the brain and behavioural similarity ratings rather than a participant specific one.

We recruited 110 participants to take part in the triplet odd-item-out task using Prolific. This sample of participants did not participate in the EEG experiment. The age data were lost for 10 participants and the gender data for 12 participants, and the demographics reported here are based on the remaining participants. Participants had a mean age of 30.07 years (SD = 9.30 years, range 20-65 years). Forty-six participants identified as female and 58 as male. The participants were healthy adults and received payment for their participation. The study was approved by the Human Ethics Committee of the University of Sydney and all participants provided informed consent before participating.

We used the same 154 food images as in the EEG experiment. Participants completed 150 trials, and the experiment took approximately 35 minutes to complete. Note that a subset of the possible combinations of triplets based on 154 items was tested for each participant. Each participant saw a unique combination of triplets, but even across participants it was not possible to test all triplet combinations. This means that a different number of observations contributes towards the dissimilarity score in each cell of the RDM, and 1.54% of cells have no observations. Hebart and colleagues (2020) recently showed that a sample of only 0.14% of possible triplet combinations contained enough information to reliably predict the entire RDM. The cells with no observations were excluded from the correlation analyses.

### 2.7. Statistical analysis

We used Bayesian statistics (Dienes, 2011; Kass & Raftery, 1995; Morey et al., 2016; Rouder et al., 2009; Wagenmakers, 2007) to assess whether there was evidence for above chance decoding accuracies or for (partial) correlations above 0. We used the Bayes Factor R package (Morey & Rouder, 2018) for all statistical analyses. We applied all statistical analyses at the group level.

For the decoding analysis, we used the same analysis for both the fixation monitoring and food categorisation tasks. We applied Bayesian t-tests for each time-point to determine the level of evidence in favour of chance decoding (null hypothesis) or above chance decoding (alternative hypothesis). We used a point null for the null hypothesis and a half-Cauchy prior for the alternative hypothesis to make the test directional (Morey & Rouder, 2011). The prior was centred around chance-level decoding (i.e. d = 0) and we used the default prior width of d = 0.707 (Jeffreys, 1998; Rouder et al., 2009; Wetzels et al., 2011). We excluded the interval between d = 0 and d = 0.5 from the prior to exclude small effect sizes, as these have been shown to occur at chance under the null hypothesis (Teichmann et al., 2022). We repeated this Bayesian t-test analysis for each time-point in the epoch.

For the representational similarity analysis, we used a Bayesian t-test for each time-point to test the (partial) correlation against 0. We applied the same Bayesian t-tests for the fixation monitoring task as well as the food categorisation task. We used the same point null as described above, and a half-Cauchy prior for the alternative hypothesis to test for positive correlations. We used the default prior width of 0.707 and excluded the interval between d = 0 and d = 0.5 from the prior for consistency with the t-test used for the decoding analysis.

Bayes Factors (BFs) above 1 show evidence in favour of the alternative hypothesis (above chance decoding or positive correlation) and BFs below 1 show evidence for the null hypothesis (chance decoding or no correlation). BF > 3 or BF < 1/3 have been interpreted as some evidence and BF > 10 or BF < 1/10 as strong evidence (Schmalz et al., 2020; Wetzels et al., 2011). Here, we use the second consecutive BF > 10 to define the onset time of above chance decoding.

## 3. RESULTS

### 3.1. The decoding of food and naturalness

To determine the time-course of the representation of visually presented food items in the brain, we decoded food vs. non-food object items for the fixation monitoring task. Participants passively viewed sequences of food and non-food object images at 6.67 Hz. Figure 4A shows the edibility decoding accuracy. There was information in the EEG signal about whether an object was food, starting around approximately 80 ms after stimulus onset with peak food decoding at approximately 196 ms. These results indicate that at approximately 80 ms, the brain encodes the distinction between food and non-food objects. To determine whether there could be a contribution from eye movements signals to these results, we repeated the decoding analysis on 56 selected posterior electrodes. This captures the signals from occipital, temporal, and parietal regions, while minimising susceptibility to contamination from eye movements picked up by frontal electrodes (Lins et al., 1993), although the contribution of eye movements cannot be fully ruled out. Figure 4A shows the food decoding accuracy for the posterior 56 electrodes. We observe the same pattern of results for the decoding based on the posterior 56 electrodes as all electrodes. There was evidence for edibility decoding from 80 ms after stimulus onset.

**Figure 4.**
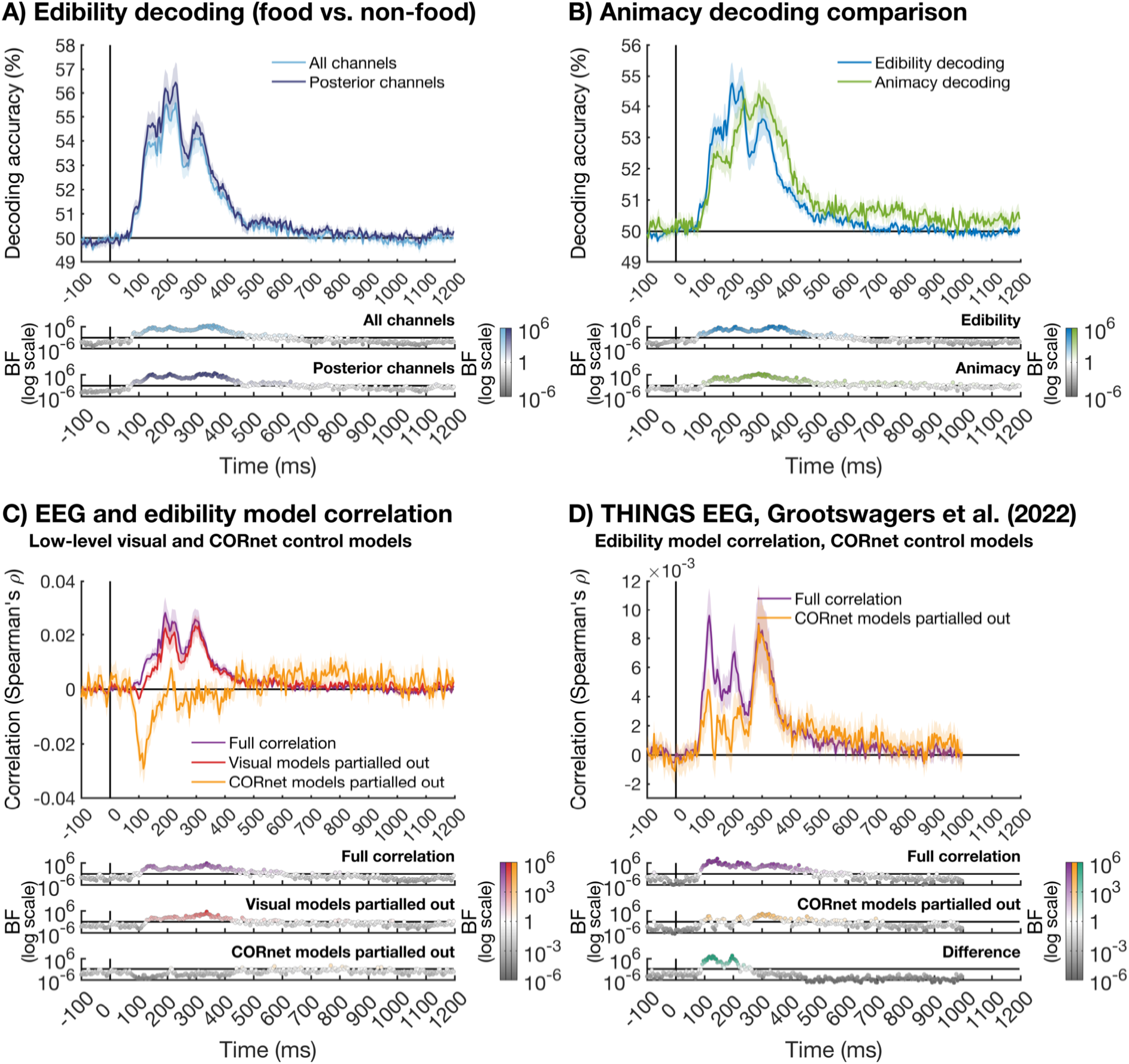
The time-course of food representation. The shaded area around the plot lines shows the standard error of the mean. The Bayes factors (BFs) are shown below the plots on a logarithmic scale. BFs > 1, plotted in the corresponding line colour, show evidence for above chance decoding and BFs < 1, plotted in grey, show evidence for chance decoding. White circles show insufficient evidence (BF close to 1). **A)** Shows the decoding accuracy of food vs. non-food object decoding over time. Theoretical chance is at 50% decoding accuracy. The light blue line shows the decoding based on all channels, whereas the dark blue line shows the decoding based on the posterior 56 channels only. **B)** shows animacy decoding in green as a comparison. We repeated the edibility decoding analysis, matching the number of training trials with the animacy analysis. Edibility decoding is shown in blue. **C)** shows the correlation between the EEG RDMs and the edibility model over time. The purple line shows the full correlation, the red line shows the partial correlation with the low-level visual control models partialled out, and the yellow line shows the correlation with the CORnet control models partialled out. The BFs are shown below the plot. **D)** shows the correlation between the EEG RDMs and the edibility model over time for the EEG dataset from Grootswagers and colleagues (2022). Following Figure 4C, the purple line shows the full correlation, and the yellow line shows the correlation with the CORnet control models partialled out. The BFs are shown below the plot, with the bottom panel showing the BFs for the difference between the full and partial correlation (teal). This is based on a one-tailed test of full correlation > partial correlation.

As a comparison, Figure 4B shows animacy decoding. Note animacy decoding was based on the non-food objects only, which means this analysis is based on fewer trials compared to the edibility decoding analysis. To match the power between analyses, we also repeated the edibility decoding, matching the number of trials with the animacy decoding analysis. There was information about edibility in the EEG data from 84 ms onwards, and about animacy from 100 ms onwards. Note that both edibility and animacy decoding could be driven, at least in part, by low-level visual differences between the groups of stimuli.

To determine whether this edibility distinction (i.e., food versus non-food objects) could be driven by low-level visual differences between the food and non-food objects, we correlated EEG RDMs with an edibility model, partialling out control models (Figure 4C). First, we controlled for the five low-level visual control models. We found evidence for a full correlation between the EEG and edibility model from 116 ms onwards. When partialling out the low-level visual control models, we found evidence for a partial correlation between the EEG and edibility model from 132 ms onwards. We also used CORnet-S layers V1, V2 and V4 as a control for low-level visual differences between food and non-food object stimuli. There was no reliable evidence for a positive correlation after controlling for the CORnet models. This means that the observed ‘edibility’ information could have been driven by visual differences between food and non-food images. This would mean that the CORnet control models provide a better control than the five low-level visual control models.

We hypothesised that the lack of a correlation between the EEG and edibility model, when using CORnet control models, could have been exacerbated by our stimulus set. The food and non-food objects were presented on a white background, making differences in shape between the two classes salient. To test this hypothesis post-hoc, we used an existing EEG dataset (Grootswagers et al., 2022) that used the THINGS stimulus set (Hebart et al., 2019), consisting of images with a natural background. We selected 277 food concepts and 280 non-food concepts. Figure 5 shows examples of selected stimuli from the THINGS stimulus set. The food concepts covered both natural and prepared items. The non-foods were chosen to cover 140 animate and 140 inanimate concepts. The animate concepts covered animals and human body parts, and the inanimate stimuli covered clothing, furniture, home décor, kitchen tools, musical instruments, office supplies, parts of a car, plants, sports equipment, tools, toys, and vehicles. A complete list of included concepts (https://doi.org/10.17605/OSF.IO/PWC4K), and all images (https://doi.org/10.17605/OSF.IO/JUM2F) can be found through the Open Science Framework. We tested for edibility information in this dataset (Figure 4D) and found edibility information from 112 ms onwards after partialling out the CORnet models. However, this information was reduced compared to the full correlation. Notably, the peak between approximately 250 ms and 380 ms was not impacted by partialling out the control models.

**Figure 5.**
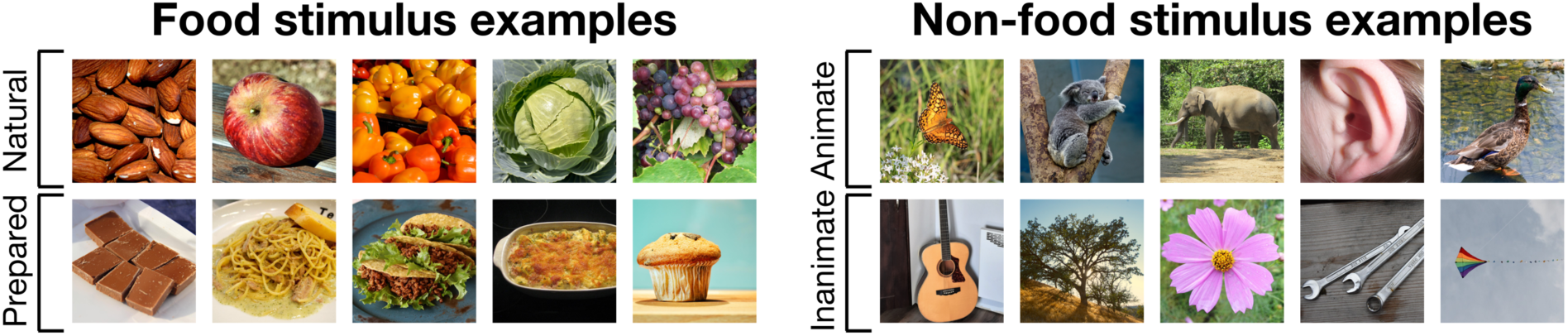
Example stimuli from the THINGS dataset (Grootswagers et al., 2022). The stimuli from the THINGS dataset were replaced with visually similar images in this Figure due to copyright. The included stimuli consisted of 277 food concepts (left) and 280 non-food concepts (right). Each concept contained 12 different images depicting the same concept. The food concepts covered both natural and prepared items, and the non-foods covered animate and inanimate items. The Open Science Framework has a complete list of included concepts (https://doi.org/10.17605/OSF.IO/PWC4K). All original images from the THINGS stimulus set (Hebart et al., 2019) can also be found via the Open Science Framework (https://doi.org/10.17605/OSF.IO/JUM2F). The visually similar matched images were obtained from publicdomainpictures.net and free-images.com.

Table 1 shows an overview of latencies for the partial correlations between the edibility model and the EEG data from the fixation monitoring task, as well as the THINGS EEG data (Grootswagers et al., 2022). Together, the results show that we need to be careful when interpreting the edibility results based on segmented images. Images with a natural background provide a better stimulus set, as these natural images reduce the salience of these visual differences. Critically, the analysis of natural images shows there is edibility information as early as 112 ms after stimulus onset.

**Table 1.**
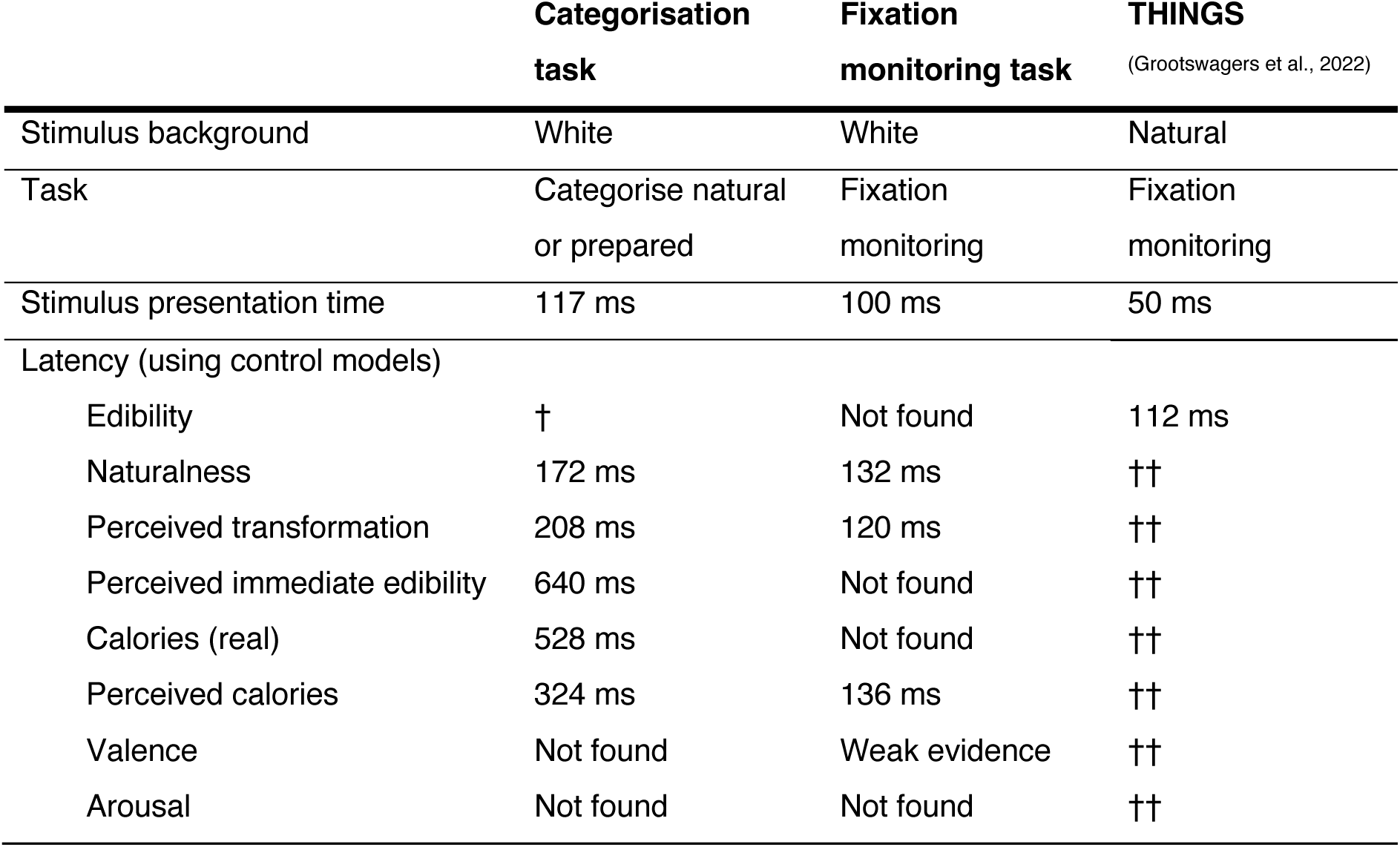
Overview of the RSA results for the categorisation task, fixation monitoring task, and the THINGS data from Grootswagers and colleagues (2022) . All analyses use control models to account for low-level visual differences between the stimulus classes. † The edibility (food vs. non-food objects) is not included, because the categorisation task did not contain non-food objects. †† The THINGS dataset was only used to test the edibility model.

To gain insight into whether the naturalness distinction between foods is represented in the brain, we decoded natural vs. prepared foods for both the food categorisation and fixation monitoring tasks. For the food categorisation task, participants actively had to think about whether a food was natural or prepared. While in the fixation monitoring task, the food stimuli were not task relevant. The decoding accuracies are shown in Figure 6. For the food categorisation task, the naturalness of the food items could be decoded approximately 116 ms after stimulus onset, with peak decoding around 436 ms. For the fixation monitoring task, the naturalness of food could be decoded approximately 108 ms after stimulus onset, with peak decoding observed around 236 ms. We observed the same pattern of results when only the posterior 56 electrodes were used in the decoding analysis, with an above chance decoding onset of naturalness information from 108 ms after stimulus onset in the food categorisation task and 104 ms after stimulus onset in the fixation monitoring task. It is possible that low-level visual differences between the natural and prepared stimuli contributed to the decoding of naturalness information. We therefore repeated the naturalness analysis using RSA, using eight models to control for these possible differences (see section 3.2). The naturalness decoding results should be interpreted together with the naturalness RSA results. Taken together, the decoding results show that whether an object is a food is represented in the brain. Using segmented images, the edibility information was driven by visual differences between food and non-food stimuli. However, when using natural images, there was information about edibility that was not driven by visual differences from 112 ms onwards. The naturalness of a food item is represented from approximately 108-116 ms after stimulus onset, although we did not control for low-level visual differences in this analysis (see section 3.2 for the control analysis). Interestingly, both types of information are represented when the food stimulus is not task relevant. However, it is still unclear whether additional features of food are represented in the brain. In the subsequent analyses, we used RSA to address this question.

**Figure 6.**
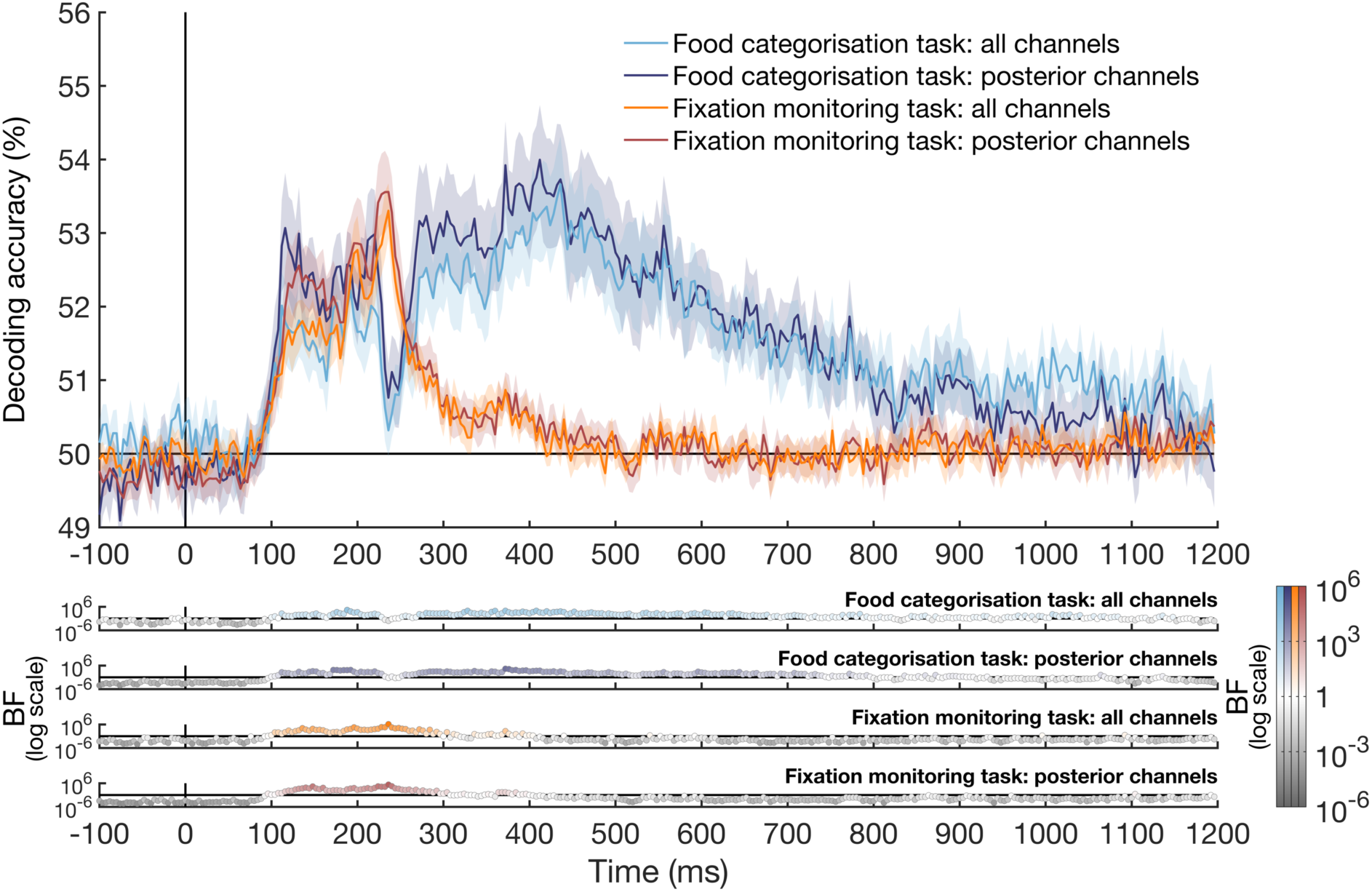
The time-course of naturalness representation in food items. The plot lines show the decoding accuracy for natural vs. prepared decoding for the food categorisation task (cool plot colours) and the fixation monitoring task (warm plot colours). The lighter coloured lines show the decoding accuracy based on all channels, whereas the darker coloured lines show the decoding accuracy based on posterior 56 channels. Plotting conventions are the same as in Figure 4. The shaded area around the plot lines indicates the standard error of the mean, and 50% decoding accuracy is theoretical chance. Bayes factors are shown on a logarithmic scale below the plot. BFs > 1, indicating evidence for above chance decoding, are shown in the corresponding plot colours. BFs < 1, indicating evidence for chance decoding, are shown in grey.

### 3.2. Representation of food properties

We used RSA to determine which dimensions of food were represented in the EEG signal at each time-point. We calculated partial correlations between the EEG RDMs for the fixation monitoring and food categorisation tasks and seven different food models: naturalness, perceived transformation, perceived immediate edibility, calories, perceived calories, valence, and arousal. Note that the food feature models were based on behavioural ratings collected by Foroni and colleagues (2013) as part of the FRIDa stimulus set. To minimise the potential contribution of low-level visual differences between the stimuli to these correlations, we controlled for eight models: three CORnet-S models and five low-level image feature control models. An overview of latencies for the partial correlations between the food models and the EEG data from both tasks can be found in table 1. Figure 7 shows the partial correlation between the food models and the EEG data from the food categorisation task, controlling for the visual models. There was evidence for a partial correlation (BF > 10) between the EEG and naturalness and perceived transformation, starting around 376 ms and 208 ms, and peaking around 444 ms and 432 ms, respectively. There was also evidence for a partial correlation between the EEG and the perceived immediate edibility model. This information emerged around 640 ms after stimulus onset, which is later compared to the onset of naturalness and perceived transformation information. There was evidence for no correlation with the *real* calorie content for most of the time-course except for a few time-points in a small window around 528 ms – 548 ms. However, there was evidence for a correlation with the *perceived* calories of the foods. Evidence for the correlation with the perceived calories emerged approximately 324 ms after stimulus onset, and the partial correlation peaked at 424 ms. There was evidence for no partial correlation between the EEG and either valence or arousal. Note that the valence and arousal models were based on ratings from a different cohort of participants. This means these models might not have been a good fit between these groups. We explore this point further in the discussion.

**Figure 7.**
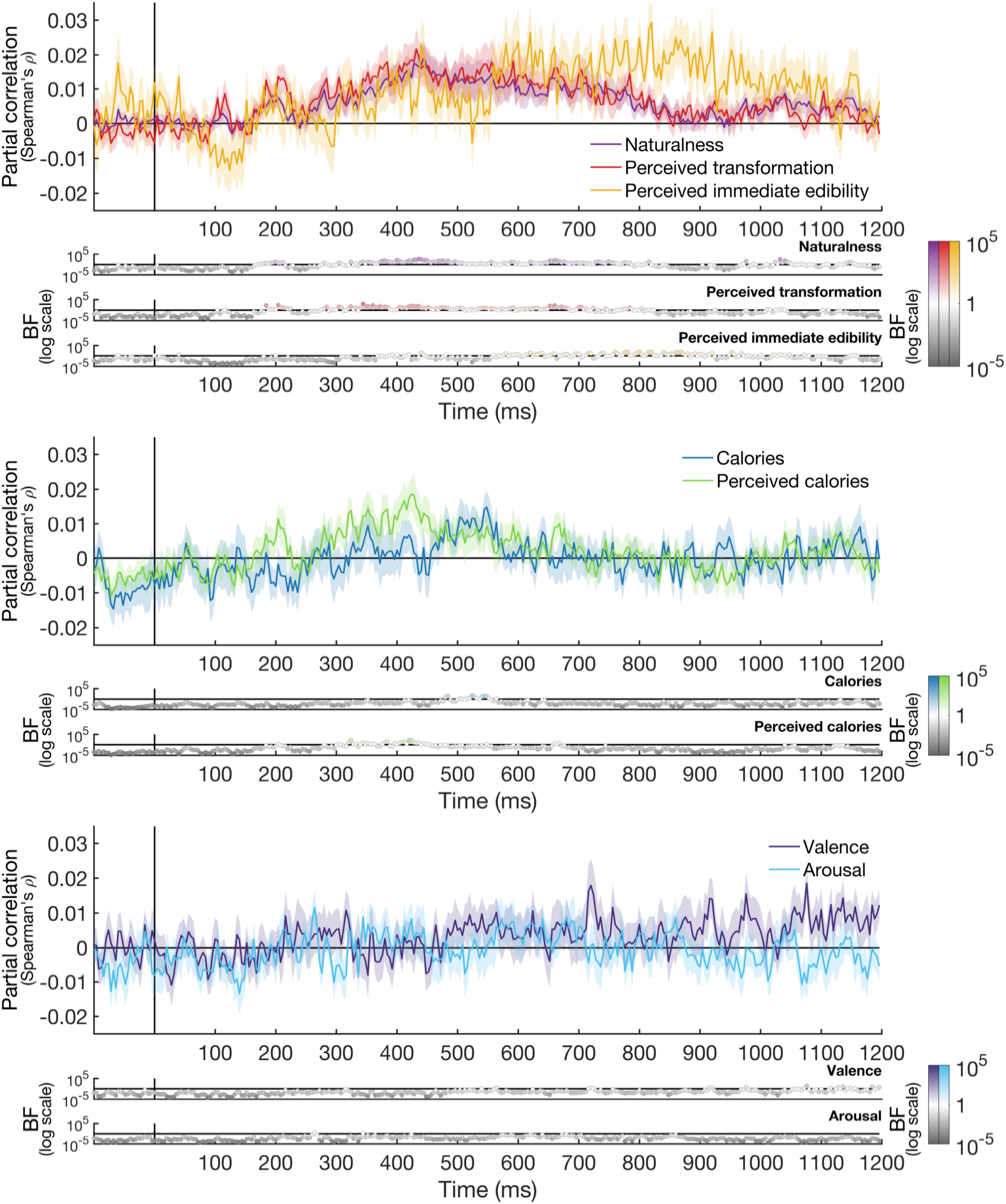
Partial correlations between the food feature models and the *food categorisation task* EEG. The five low-level visual models and three CORnet control models were partialled out. Plotting conventions are the same for all three panels. The top panel shows the partial EEG correlations with the naturalness model (purple), perceived transformation model (red), and perceived immediate edibility model (yellow). The shaded areas around the plot lines show the standard error of the mean. The Bayes Factors are shown below the plot on a logarithmic scale. BFs > 1 are shown in the corresponding plot colour: purple for the naturalness model (top), red for the perceived transformation model (middle), and yellow for the perceived immediate edibility model (bottom). They indicate evidence for a partial correlation. BFs < 1 are shown in grey and indicate evidence for chance decoding. The middle panel shows the partial correlations between the EEG and the real calories model (blue) and perceived calories model (green). The Bayes Factors are shown below for the real calories model (top) and perceived calories model (bottom). The bottom panel shows the partial EEG correlation with the valence model (purple) and the arousal model (light blue). The corresponding Bayes factors are plotted below for the valence model (top) and the arousal model (bottom).

The data show that the naturalness, perceived transformation, and perceived calorie models can explain part of the variance in the EEG data. However, it is unclear whether these findings are driven by the task, as participants were asked to report whether the food was natural or prepared, making the naturalness and perceived transformation dimensions task relevant. To determine whether the findings hold when the foods are not task relevant, and presented at a fast rate, we correlated the EEG for the fixation monitoring task with the food models, again partialling out the eight visual control models (Figure 8). Overall, the pattern of results is similar to the active food categorisation results. There was a correlation between the EEG and 1) the naturalness model and 2) the perceived transformation model from approximately 180 ms and 220 ms after stimulus onset respectively, both peaking at 232 ms. Unlike the results for the food categorisation task, there was evidence for no partial correlation between the EEG and perceived immediate edibility for most of the time-course. As in the food categorisation results, the fixation monitoring results showed evidence for no partial correlation between the EEG data and the *real* calorie content, but there was evidence for a partial correlation between the EEG and the *perceived* calorie model. Evidence for the latter emerged approximately 256 ms after stimulus onset. Finally, the valence model showed no evidence for a partial correlation with EEG for most time-points, with some evidence in favour of a partial correlation (BF > 3) for a few time-points between 332 ms and 368 ms. There was evidence for no partial correlation between the EEG data and the arousal model. Taken together the results show that the naturalness, perceived transformation, and perceived calorie models explain part of the EEG variance, whereas the real calories, valence and arousal models have little to no explanatory power. This pattern of results holds for the fixation monitoring task, where the stimuli are not task relevant and presented at a rapid presentation rate (6.67 Hz). There was a late correlation between the EEG and perceived immediate edibility model, but this was only observed for the food categorisation task.

**Figure 8.**
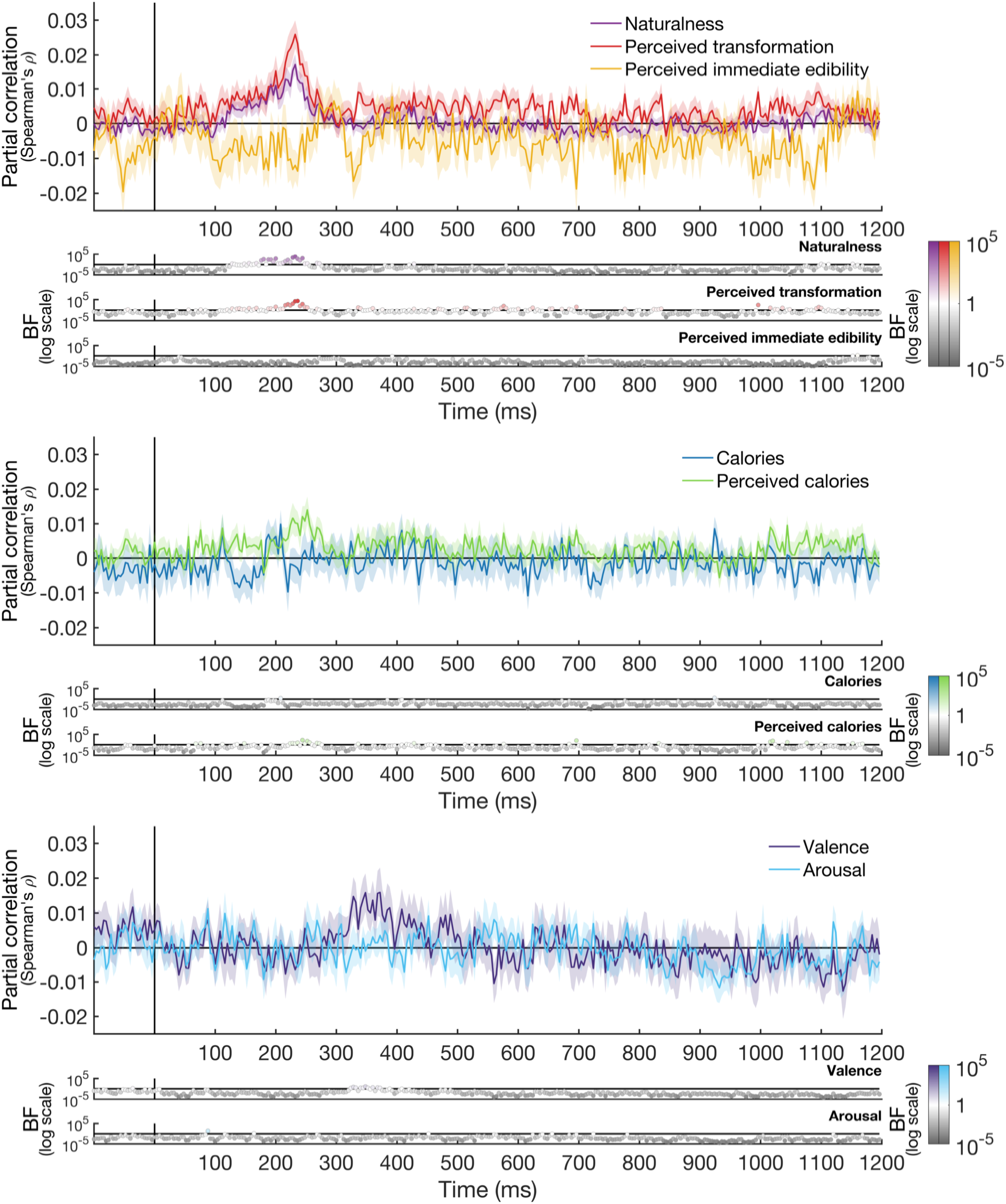
Partial correlations between the food feature models and the *fixation monitoring task* EEG. All plotting conventions are the same as in Figure 7.

We used partial correlations to determine whether the correlated food feature models provide unique contributions to the correlations between the models and the EEG RDMs. Specifically, we investigated the correlation between the EEG and 1) naturalness, 2) perceived transformation, and 3) perceived calories, partialling out one or more correlated models. We did this separately for the categorisation task (Figure 9A) and the fixation monitoring task (Figure 9B). The naturalness and perceived transformation models share most of their variance. No time-points with BFs over 10 remain when partialling out one model from the correlation between the other model and the EEG in either task. The exception is the correlation between perceived transformation and the EEG for the fixation monitoring task, where a few time-points have BFs over 10 when partialling out naturalness. The shared explanatory power between the naturalness and perceived transformation models is expected, as both models are conceptually similar. The ‘perceived transformation’ is a continuous and more subjective version of the ‘naturalness’ model. We also investigated the unique contribution of the perceived calorie model. Interestingly, the perceived calorie model provides very little unique contribution over the naturalness and perceived transformation models, with only a few time-points showing BFs above 10 for both tasks when either model is partialled out. However, the naturalness and perceived transformation models do have a unique contribution above the perceived calorie model, where clusters of time-points with BFs over 10 remain when the perceived calorie model is partialled out. In addition, partialling out the real calories from the correlation between the EEG and perceived calorie model does not appear to diminish the correlation. This is expected, as although the real and perceived calorie models are correlated, the real calorie model does not have explanatory power regarding the EEG data. Taken together, these results show that the naturalness and perceived transformation model explain shared variance in the EEG signal. They can explain EEG variance over and above the perceived calorie model, but the perceived calorie model itself explains very little unique variance in the EEG.

### 3.3. Behavioural human food discrimination model

We used RSA to study what features humans use to discriminate food stimuli. We used behavioural responses from an odd-one-out triplet experiment, completed by a different group of participants, to make a behavioural RDM (Figure 10A). Importantly, the feature that was used to make an odd-one-out decision was not constrained, and participants could use any feature they wished to make this decision. To gain insight into the features that contribute to participant’s odd-one-out decisions, we correlated the behavioural RDM with the seven food feature model RDMs (Figure 2B) and the five low-level image feature control model RDMs (Figure 2C). Figure 10B shows the correlations between the behaviour and the food/visual models. All low-level image feature models had relatively low correlations with the behavioural RDM, suggesting that participants did not often rely on visual differences between stimuli to make their odd-one-out decision. The highest correlations were between the behavioural RDM and perceived transformation, naturalness, and perceived calories. This suggests that participants often rely on one or more of these features to inform their odd-one-out decision. Note, these three food models are correlated (see Figure 2D and 2E), which means it is difficult to distinguish between these features in terms of the driver behind behavioural odd-one-out decisions.

**Figure 9.**
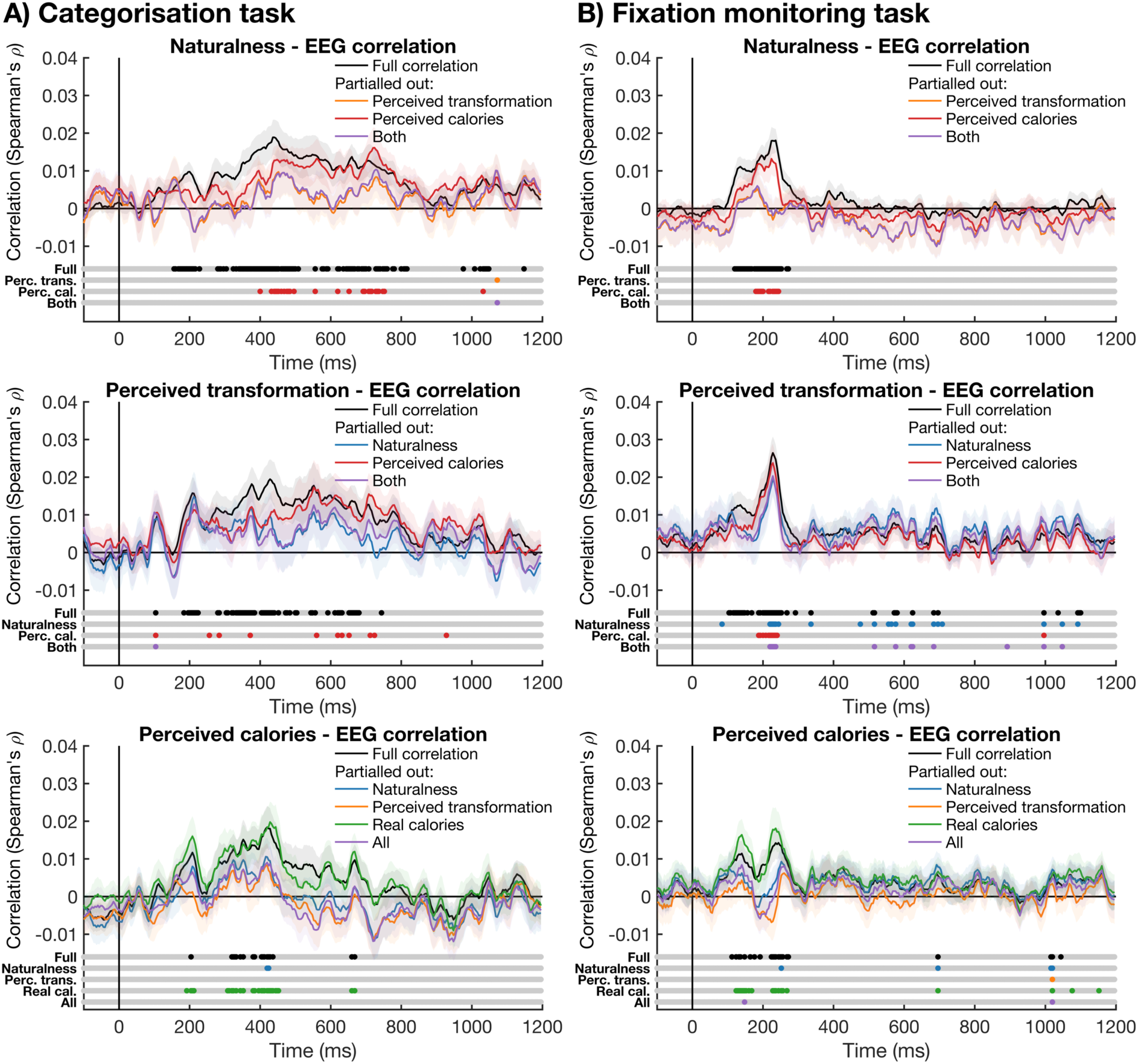
Unique model contributions to the correlation with EEG. **A)** shows the model contributions for the categorisation task (temporally smoothed with a kernel of 5 time-points for visibility). The top panel shows the full correlation between the EEG RDM and the naturalness model (black), as well as three partial correlations, partialling out 1) the perceived transformation model (yellow), 2) the perceived calorie model (red), and 3) both (purple). The middle panel shows the full correlation with the EEG and the perceived transformation model (black), and three partial correlations, partialling out 1) the naturalness model (blue), 2) the perceived calorie model (red), and 3) both (purple). The bottom panel shows the full EEG correlation with the perceived calorie model (black) and four partial correlations, partialling out 1) the naturalness model (blue), 2) the perceived transformation model (yellow), 3) the real calorie model (green), and 4) all these models (purple). The Bayes factors are shown below each plot (no temporal smoothing was applied). BFs > 10 are shown in the plot colour and BFs < 10 are shown in grey. **B)** shows the model contributions for the fixation monitoring task. All plotting conventions are the same as for Figure 9A.

**Figure 10.**
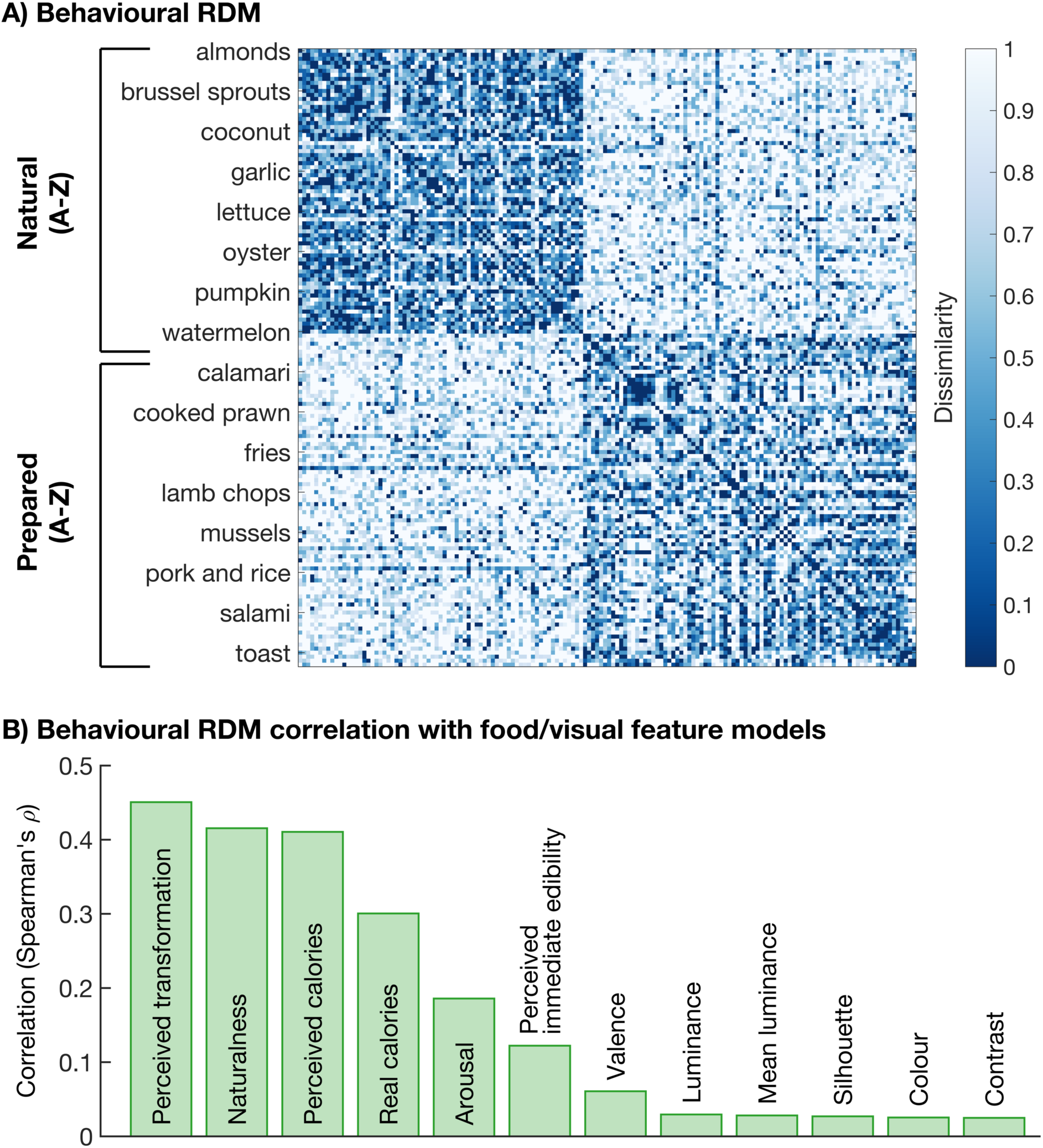
Behavioural human food discrimination model. **A)** Shows the behavioural RDM, based on the odd-one-out triplet task. Each point in the 154 by 154 RDM represents the dissimilarity between two food stimuli. The food items are ordered as follows: all natural foods in alphabetical order, followed by all prepared foods in alphabetical order. **B)** Shows the correlations between the behavioural RDM and the food/visual feature model RDMs. We included seven food feature models (see Figure 2B) and five low-level image feature models (see Figure 2C).

### 3.4. The link between the neural representation of food features and behaviour

The results above elucidate the emerging representation of visually presented food stimuli. To determine whether this information is represented in a way that is accessible to the brain, and used to guide behaviour, we aimed to establish a link between the neural representations and behaviour obtained from two different groups of participants. To this end, we correlated the EEG with the behavioural RDM from the triplet data. Figure 11 shows the time-course of the correlation between the EEG for the two tasks and behaviour. There was a correlation between the EEG for the food categorisation task, where participants indicated whether a food was natural or prepared, and the behavioural RDM from approximately 184 ms onwards. Interestingly, there was also a correlation between the EEG data for the fixation monitoring task and the behavioural RDM, even though the food was not task relevant in the EEG task. This correlation emerged around 136 ms after stimulus onset, with a clear peak at 232 ms after stimulus onset. Together, this pattern of results suggests that the features that guide behavioural similarity judgments are represented in the neural signal from approximately 136- 184 ms onwards, for both tasks.

**Figure 11.**
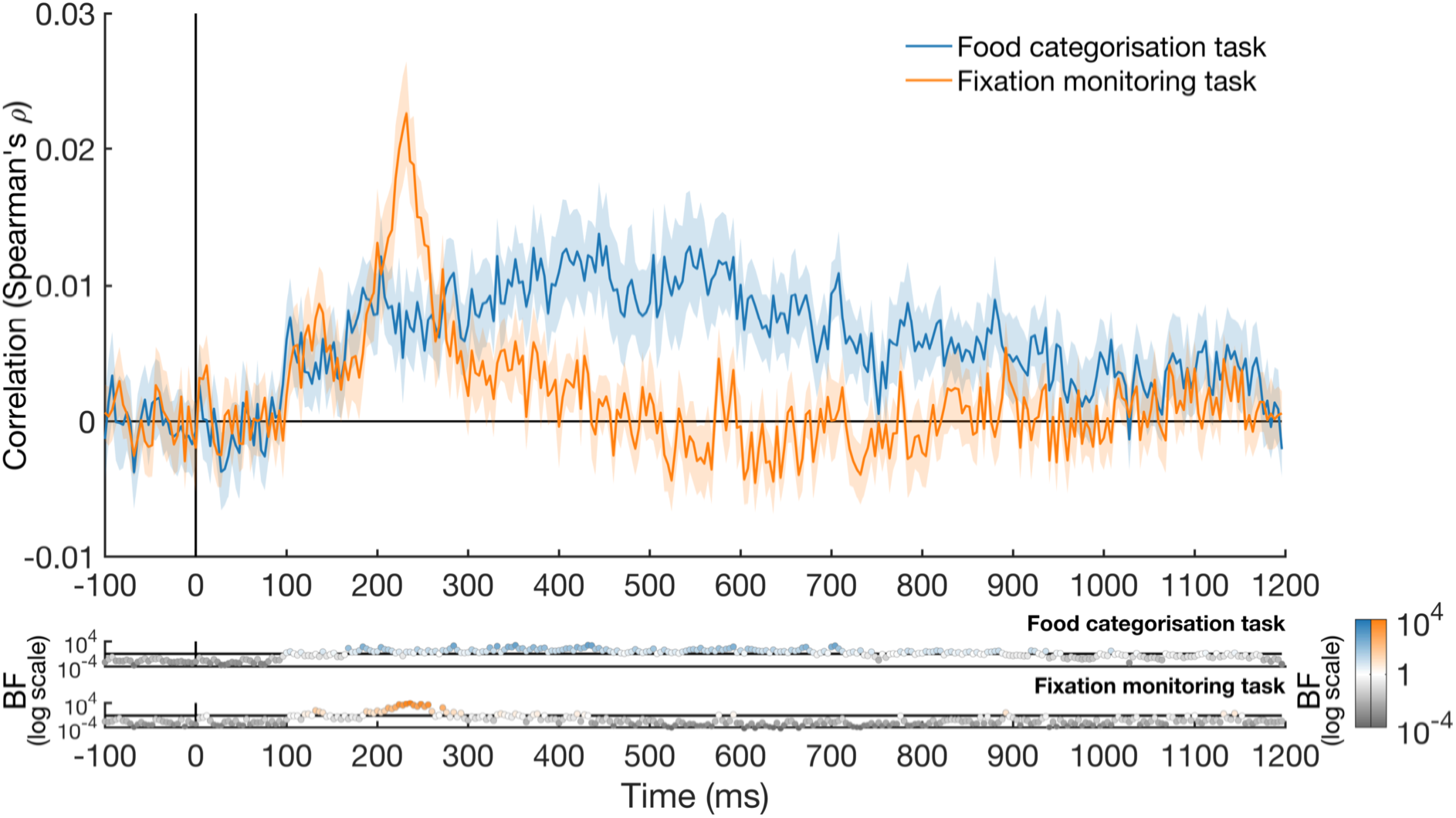
The correlation between the behavioural model RDM and the EEG for the two different tasks. The correlation with the EEG for the food categorisation task is shown in blue and the correlation with the EEG for the fixation monitoring (passive viewing) task is shown in orange. The shaded areas around the plot lines show the standard error of the mean. Bayes factors are shown below the plot. BFs > 1, are shown in blue for the food categorisation task (top) and in orange for the fixation monitoring task (bottom). BFs < 1 are shown in grey.

## 4. DISCUSSION

In this study, we used time-resolved multivariate analyses to explore the brain processing dynamics of visual food stimuli and asked whether the food item must be task relevant for the brain to represent food stimuli and features (e.g., perceived calories). When using segmented images, the distinction between food and non-food items in the brain was driven by visual differences between stimuli. However, when using natural images, there was information about edibility from 112 ms after presentation. The neural signal contained information about food naturalness, the level of transformation, and perceived caloric content, for both tasks, whereas perceived immediate edibility was only represented when participants engaged in the food categorisation task. In addition, the recorded brain activity correlated with the behavioural responses of a different group of participants in an odd-item-out task.

The finding that the brain represents edibility information is in line with previous work using fMRI (Avery et al., 2021; Chen et al., 2016; Jain et al., 2023; Khosla et al., 2022; Killgore et al., 2003; Rothemund et al., 2007; Simmons et al., 2005) and MEG (Stingl et al., 2010; Tsourides et al., 2016). We found that edibility information was represented from 80 ms onwards using segmented stimuli, in line with previous work (Tsourides et al., 2016). However, using RSA with CORnet control models, we found that there was no longer evidence for a correlation between the EEG and edibility model (for segmented stimuli) or a reduction in the correlation before 250 ms using THINGS natural images (Grootswagers et al., 2022; Hebart et al., 2019). This means that care must be taken when interpreting the onset times based on segmented stimuli. Importantly, using natural images, we find information of edibility information emerges after 112 ms, indicating the brain rapidly distinguishes food from non-food stimuli. Notably, the onset time of edibility information occurs at a similar time as information about subjective relevance and wanting ratings (Turner et al., 2017), which was present in the 100 ms to 150 ms window after stimulus onset.

Our results also show that the brain represents whether food is natural or prepared. This distinction emerged approximately 180-376 ms after stimulus onset and was observed even when controlling for low-level visual information. This finding is consistent with previous EEG work that found differences in amplitude in response to natural compared to prepared foods (Coricelli et al., 2019; Pergola et al., 2017). Our work builds on previous findings by showing that the brain represents information about food naturalness. Finally, this information was observed even when the stimuli were presented at a fast rate (6.67 Hz) and the food was not task relevant. These findings are consistent with rapid processing by the brain of naturalness information in visually presented food and suggest that the level of transformation is an important dimension along which the brain organises food representations.

In addition to representing food naturalness, our results show that the brain represents the perceived energetic/caloric content of food. We observed partial correlations between the EEG and perceived calories, emerging around 256 ms after stimulus onset for the fixation monitoring task. This finding is in line with previous work from fMRI (Killgore et al., 2003; Mengotti et al., 2019) and EEG (Toepel et al., 2009), and suggests that this information could be represented automatically. Our study adds to previous work by showing that the brain represents information about the energetic/caloric content of the food. Interestingly, the EEG did not show reliable evidence for a correlation with the *real* caloric content of the food, except for a small window around 528 ms – 548 ms in the stimulus categorisation task. As the perceived calorie model is not based on the participants from the EEG task, it is therefore not personal biases in calorie perception, but rather generalisable ideas about energetic value of foods that are represented in the neural signal.

We also found evidence for information about the perceived immediate edibility of the food in the food categorisation task, although this information emerged much later compared to information about naturalness. We did not find immediate edibility information for the fixation monitoring task, which could mean that the coding of perceived immediate edibility is not automatic and is only present when attention is oriented towards the food item. Although the food is task relevant in the food categorisation task, the ‘perceived immediate edibility’ dimension itself is not relevant for the task, as participants are asked to rate whether the food is natural or prepared. The naturalness and perceived immediate edibility dimensions are not strongly correlated (r = 0.19), suggesting that attending to an unrelated feature of a food item could be sufficient for the coding of perceived immediate edibility to occur. However, note that the timing of the fixation monitoring task was also faster, which means there could have been earlier masking in this task (Robinson et al., 2019).

Finally, our results showed that the brain did not reliably represent general valence or arousal. This finding seems inconsistent with Schubert and colleagues (2021), who found that personal subjective tastiness (valence) could be decoded from 530-740 ms onwards, depending on the task. There are a few possible explanations for this apparent difference. First, the behavioural valence and arousal ratings in our task were based on a different cohort of participants, and therefore represent general preferences across the population rather than personal preference. It is possible that these subjective ratings did not transfer across participants, and only personal preference is reliably coded in the brain. A larger range of food stimuli, including low valence / high arousal foods such as rotten foods, might tap into universal representations of valence and arousal. Alternatively, the lack of this information could also be driven by the internal state of the participant. Participants in the study by Schubert and colleagues (2021) were asked to fast for 4 hours before the study, whereas no specific instructions about fasting were given in this study. It is possible that temporary states, such as hunger, as well as longer-term differences, such as diet and body mass index, can mediate the coding of different food features. Future work can further investigate the degree of universality and individuality of the representation of food features in the brain.

We have shown that the brain represents various food features, such as edibility, naturalness, and perceived caloric content. To directly test whether this information is used by the brain to guide behaviour, we correlated the EEG with a behavioural RDM, based on a triplet odd-item-out task done by a different group of participants. Our results showed that there was a correlation between brain and behaviour from 136-184 ms onwards. This means that the EEG signal from that time and later could predict the behavioural response of a different group of participants. This correlation was found for both EEG tasks, suggesting that the information about foods is used to guide similarity judgments about the food. Interestingly, the features that were used most by participants to make an odd-one-out decision were naturalness/perceived transformation and perceived calories. These are also the features that are represented in the neural signal. In addition, because the EEG data was obtained from a different group of participants compared to the behavioural data, this suggests a degree of universality in the features that guide food similarity decisions. This adds further evidence to the importance of naturalness and energetic content in the perception of food.

The fixation monitoring task and food categorisation tasks were used to provide insight into the passive and active processing of visual food stimuli respectively. The experiment is not designed to directly compare the coding food information between the two different tasks. However, visual inspection shows a clear pattern in naturalness coding. There is information about naturalness between approximately 100 and 300 ms after stimulus onset, regardless of the task. This suggests that the early stage of food stimulus processing in the brain could be automatic and therefore unaffected by attention. However, note that the fixation monitoring task was not difficult, which means additional cognitive resources may have been available to participants to process features of the presented food. The naturalness information was only sustained over time in the food categorisation task. This could reflect the active representation of the relevant feature in short term memory and/or the decision the participant makes about this stimulus. Alternatively, there could have been masking due to the fast nature of the fixation monitoring task (Robinson et al., 2019). The prolonged coding of the attended stimulus is in line with previous attention work that showed initial representation of attended and unattended features, with extended representation of the attended feature only (Goddard et al., 2022; Grootswagers et al., 2021; Moerel et al., 2021, 2022).

The findings from this study elucidate the time-course with which different food dimensions are represented in the neural signal. Information about the naturalness and perceived caloric content of the food are represented rapidly in the brain. This suggests that these features are key organising principles in the representational structure of foods.

## 5. DATA AND CODE AVAILABILITY

The analysis code and figures can be found via the Open Science Framework (https://doi.org/10.17605/OSF.IO/PWC4K), and the dataset can be found via OpenNeuro (https://doi.org/10.18112/openneuro.ds004995.v1.0.2).

## 6. ACKNOWLEDGMENTS

This work was supported by Australian Research Council (ARC) Discovery Projects awarded to TAC (DP160101300 and DP200101787). We would like to acknowledge the University of Sydney HPC service for providing High Performance Computing resources.

